# Combined Transcriptomics and Proteomics Forecast Analysis for Potential Genes Regulating the Columbian Plumage Color in Chickens

**DOI:** 10.1101/512202

**Authors:** XinLei Wang, Donghua Li, SuFang Song, YanHua Zhang, YuanFang Li, Xiangnan Wang, Danli Liu, Chenxi Zhang, Yanfang Cao, Yawei Fu, RuiLi Han, WenTing Li, Xiaojun Liu, Guirong Sun, GuoXi Li, Yadong Tian, Zhuanjian Li, Xiangtao Kang

**Affiliations:** College of Animal Science and Veterinary Medicine, Henan Agricultural University, Zhengzhou, 450002, Henan, China; College of Animal Science and Technology, Henan University of Animal Husbandry and Economy, Zhengzhou, 450046, Henan, China

**Keywords:** “Yufen I” H line, Columbian plume, feather follicles, transcriptomic, proteomic

## Abstract

**Background:** Coloration is one of the most recognizable characteristics in chickens, and clarifying the coloration mechanisms will help us understand feather color formation. “Yufen I” is an commercial egg-laying chicken breed in China, that was developed by a three-line cross using lines H, N and D. Columbian plumage is a typical feather character of the “Yufen I” H line. To elucidate the molecular mechanism underlying pigmentation of Columbian plumage, this study utilizes the technology of high-throughput sequencing to compare the transcriptome and proteome differences in different feather follicular tissue, including the dorsal neck with black and white striped feather follicles (Group A) and the ventral neck with white feather follicles (Group B) in the “Yufen I” H line.

**Results:** In this study, we identified a total of 21,306 genes and 5203 proteins in chicken feather follicles. Among these, 209 genes and 382 proteins were differentially expressed in two different locations, Group A and Group B, respectively. A total of 8 differentially expressed genes (DEGs) and 9 differentially expressed proteins (DEPs) were found to be involved in the melanogenesis pathway. Besides, a specifically expressed *MED23* gene and a differential expressed GNAQ protein were involved in melanin synthesis. Kyoto Encyclopedia of Genes and Genomes (KEGG) analysis mapped 190 DEGs and 322 DEPs, to 175 and 242 pathways, respectively, and there were 166 pathways correlated with both DEGs and DEPs. 49 DEPs/DEGs overlapped and were enriched for 12 pathways. Transcriptomic and proteomic analyses revealed that the following pathways were activated: melanogenesis, cardiomycete adrenergic, calcium and the cGMP-PKG. The expression of DEGs was validated by real-time quantitative polymerase chain reaction (qRT-PCR) that was similar to that of RNA-seq. In addition, we found that *MED23, FZD10, WNT7B* and *WNT11* genes expression peaked at approximately 8 weeks in the “Yufen I” H line, which is consistent with the molting cycle. As both the groups showed significant differences in terms of expression of the genes studied, this study opens up avenues for study in the future to assess their exact function in color of plumage.

**Conclusion:** These common DEGs and DEPs were enriched in the melanogenesis pathway. The *MED23* and GNAQ were also reported to have a crucial part synthesis of melanin. In addition, this study is the first to reveal variations in gene and protein in the “Yufen I” H line during Columbian feather color development, and discover principal genes and proteins that would aid in the functional genomics studies in future. The results of the present study provide a significant conceptual basis for the “Yufen I” H line future breeding schemes and provide a basis for research on the mechanisms of feather pigmentation.

## Introduction

“Yufen I” is an egg-laying chicken breed in China, which was developed by a three-line cross using line H as the first male parent, line N as maternal grandparent and line D as the final male parent. As authorized by National Commission on Livestock and Poultry Genetic Resources in 2015, this breed has been bred true for at least six generations. However, a closed breeding method was used to develop the Line H by crossing the barred plumaged-original Gushi chicken with an egg-laying grandparent line C, the brown-shelled Babcock B-380. Line H is a fast plumage line and is characterized by early maturity, high egg production and the Columbian plumage pattern. Columbian plumage is a character of feather color in the “ Yufen I” H line in which the dorsal neck, tail and apex of the wing feathers have black and white stripes, and other feathers are white. This pattern was also regarded as the main pigmentation character of the “Yufen I” H line.

Complex feather coloration is likely coordinated through multiple genes that regulate diverse mechanisms. Feather color in chickens is a result of the melanin produced by the melanocytes of the feather follicles. Feathers and feather follicles are ideal tissues to explore the genetic mechanism and complexity of color patterns in birds. As a derivative of chicken skin, the feather follicles give rise to the feathers, and are capable of self-renewal, and their proliferation and differentiation result in feather formation [1–3]. In adult feather follicles, plumage pigmentation is mainly dependent on the interaction between feather follicle melanocytes and dermal papilla fibroblasts. Pigmentation activity occurs only during the growth period of feather follicles, and the transfer of melanin and pigment to keratinocytes depends on the activity of melanin precursors. The melanin or pigment is transferred to the skin and feather follicles through the regulation of signal transduction pathways [4,5]. Studies have revealed that many growth factors and receptors coordinate genes and that the environment and signaling pathways play an extremely important role during feather growth. To date, some of the confirmed pathways involved in pigmentation include cAMP pathway [6], SCF-KIT pathway [7], PI3K-Akt pathway [8], BMP signaling pathway [9], Notch pathway [10], ERK pathway [11], CREB/MITF/tyrosinase pathway [12], MCIR/(Gs-AC)/PKA pathway [13–15], Wnt/β-catenin pathway [16,17] and MAPK pathway [18,19].

Uniformity in the appearance of birds is essential in the poultry industry [20]. Although plumage color is easily observed, the genetics behind feather pigmentation is governed by both qualitative and quantitative features [21]. Research reveals that the ratio of pheomelanin and eumelanin pigmentation regulates the feather color in chickens. Eumelanin pigmentation requires that the melanoblasts migrate to the epidermis from the neural crest to ultimately reach the developing feather follicles. Another requirement is the proportion of the pigment subject to control by genes [22,23]. Melanosomes are responsible for synthesizing these pigments. These organelles are granule-like and are develop within melanoblasts which go through several steps of differentiation to form melanocytes. The preliminary steps for melanin production include the appearance of the neural crest, determination of melanoblast, migration, proliferation and differentiation [24–26]. Besides this, melanogenesis regulation after the melanocytes migrate into feather follicles is another important step. The variation in plumage may be caused by any mutations in respective genes and changes in molecules (receptors of transcription factor on cells, structural proteins, enzymes and growth factors) involved in the aforementioned process [14,27,28]. Several studies, until now have focused on genes such as *MC1R, ASIP, TYR, SLC24A5, KITLG, MITF* and *DCT* that play a role in the melanin proportion synthesis [29–33].

Many genes and pathways have been shown to be associated with pigmentation. However, research on the genetics of Columbian plumage in chickens is lacking. In this experiment, we collected the feather follicles in two parts: the dorsal feather follicles of the neck with black and white-striped feathers (Group A) and the ventral feather follicles of the neck with white feathers (Group B). This study aimed to discover the differentially expressed genes (DEGs), differentially expressed proteins (DEPs) and Kyoto Encyclopaedia of Genes and Genomes (KEGG) pathways by transcriptomic (RNA-seq) and proteomic (iTRAQ) analyses and to forecast potential candidate genes of Columbian plumage in order to determine which genes and pathways affected feather coloration.

## Materials and methods

### Disclosure of ethics

Experimental animals were kept as rules in National standards for the environment and facilities of experimental animals of China (GB14925-2010). All the chickens were healthy with a coop of 0.12m^2^ and 0.4m high for each. Adequate and clean drinking water and feed were provided. All the protocols for animal experiment received approval from the Institutional Animal Care and Use Committee (IACUC) of Henan Agricultural University and from Henan Agricultural University’s Animal Care Committee, College of Animal Science and Veterinary Medicine, China (Permit Number: 17-0322).

### Animals for experiments and tissue sources

The Animal Center of Henan Agricultural University provided us with chickens of the “Yufen I” H line breed for this study. Three 22-week-old female chickens were anaesthetized with “Su-Mian-Xin” (one type of anesthetic, Shengda Animal Pharmaceutical Co., Ltd, Dunhua, China), which is diluted with saline in a volume ratio of 1:2. Finally, the chicken was anesthetized by i.v. injection in the wing vein with the concentration of 0.2mL/kg for about 20 minutes and then the neck feather was plucked. New feathers emerged in the skin after two weeks. Then three chickens involved in this study were placed in an airtight box and humanely sacrificed by inhaling carbon dioxide in order to reduce their suffering. Six samples of tissues including feather follicle and the circumjacent skin closely around the rachis of the feathers were collected from the dorsal and ventral areas, the locations at which black and white-striped feathers and white feathers occur, respectively (Fig 1). Three replicates were collected for each group (A1 to A3 for Group A and B1 to B3 for Group B). Approximately 20~30 feather follicles were placed into 2-mL tubes, and then the tube was sealed, dipped in liquid nitrogen to freeze quickly and −80 °C storage until isolation and sequencing of RNA and qRT-PCR analysis [34].

**Fig 1.**
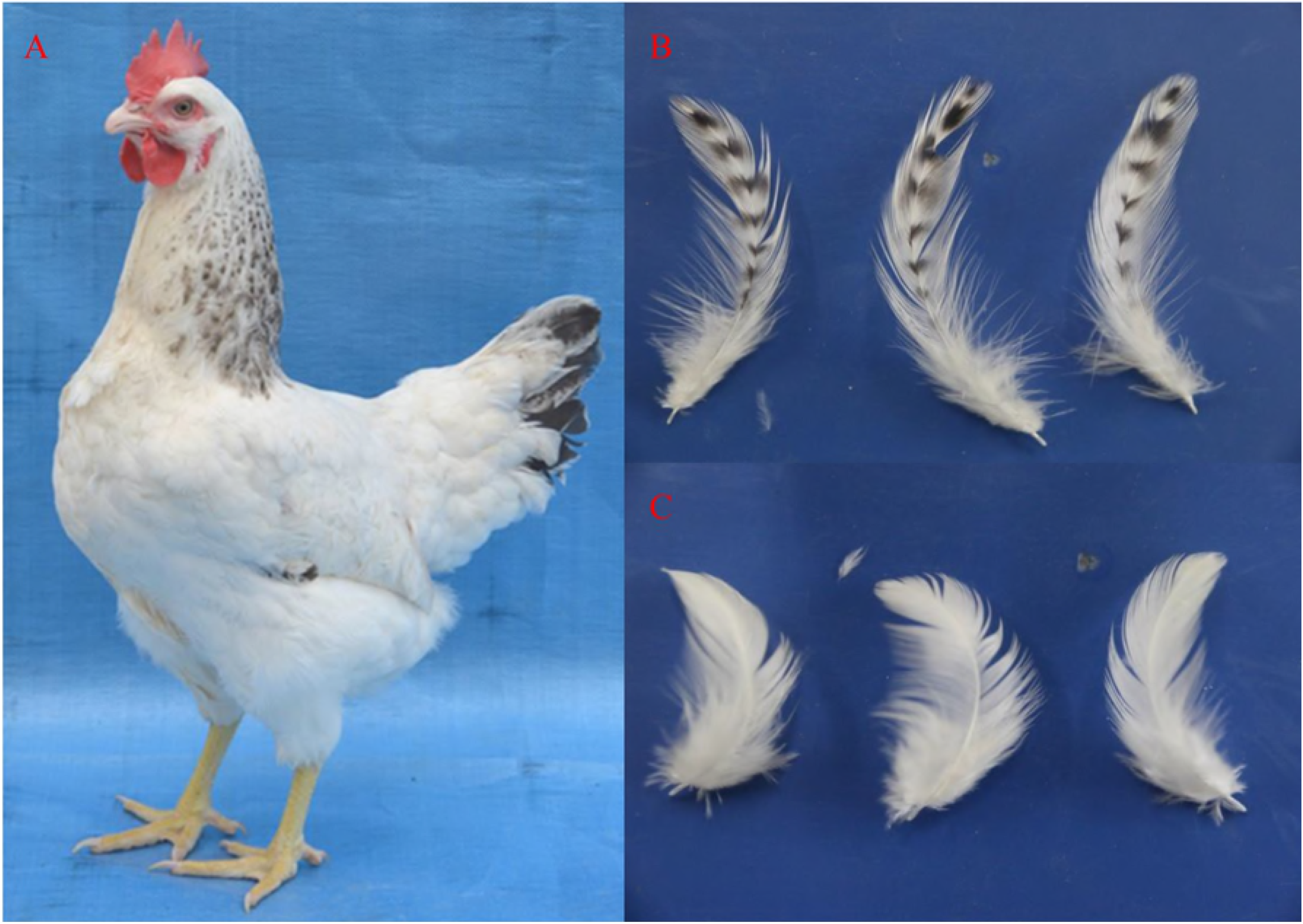
“Yufen I” H line chicken and the feather bulbs used in this study. (A). “Yufen I” H line chicken, (B). the black and white striped-feather in dorsal neck, (C). the white feather in ventral neck.

### Extraction of total RNA and RNA-seq

Six frozen samples corresponding to two the different feather follicle areas (Group A and Group B) were selected for isolating total RNA. The feather follicles of the “Yufen I” H line were used to extract total RNA using TRIzol reagent (Invitrogen, USA) and used in library construction. The assessment of contamination and degradation in RNA were done with standard denaturing agarose gel electrophoresis. The RNA integrity, concentration and purity were evaluated on an Agilent RNA6000 Nano Chip in Reagent Port 1 of the Bioanalyzer Agilent 2100 (Agilent Technologies, CA, USA). Sequencing of the libraries was done on the Illumina HiSeq 4000 platform by BGI Co., Ltd. Deposition of data was done in the NCBI Sequence Read Archive under Accession SRR7973871. For processing the raw data, its FASTQ format were first processed through SOAPnuke (v1.5.2), further, reads carrying adapters, ploy-N sequences and having low quality were then deleted from the raw data to obtain clean data. Additionally, we calculated the Q20 and Q30 at error rates of 1% and 0.1% respectively of the clean data. To ascertain if resequencing was required, quality control (QC) for alignment was carried out. This high quality data was used for the analyses performed downstream.

### Transcriptomic data processing

After read filtering, we used HISAT (v0.1.6-beta) to perform genome mapping. For mapping the reads RNA-seq, a spliced alignment program HISAT is fast and sensitive with an accuracy that is equal to or better than other methods. The spliced mapping algorithm of HISAT has been applied for mapping genome on the preprocessed reads [35,36]. Reads that passed the QC test were aligned to a reference genome assembly of chicken (*Gallus gallus 5.0, https://www.ncbi.nlm.nih.gov/assembly/GCF_000002315.4/*) from NCBI. The measurement of abundance of expression for each assembled transcript was done using the Fragments per Kilobase of exon model per Million mapped reads (FPKM) values.

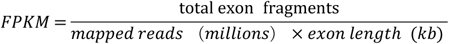

For the two groups, the analysis of differential expression, the levels of gene and transcript expression were determined using RSEM software (V1.2.12). For each pair of samples, the DEGs were screened using a model derived from PossionDis software [37]. In this paper, “false discovery rate (FDR) ≤ 0.001 and the fold change ≥ 2 (absolute value of log2Ratio ≥ 1)” were set as the significant threshold value to ascertain the differences between the gene expression. The phyper function of R software was used to identify enriched KEGG (http://www.kegg.jp/) pathways (p ≤ 0.05) and Gene Ontology (GO: http://www.geneontology.org), respectively. Additionally, for identifying the major metabolic and signal transduction pathways, KEGG enrichment analysis was done on the DEGs. Likewise, the GO enrichment analysis to ascertain the main molecular functions, cellular components and biological processes associated with DEGs.

### RNA validation through qRT-PCR

For validating the RNA-seq data, qRT-PCR was used and seven DEGs were selected for the analysis. The same samples used for RNA sequencing were used for reverse transcription and synthesis of cDNA with PrimeScript RT Reagent Kit with gDNA Eraser following the instructions of the manufacturer (TaKaRa, Dalian, China). For designing the primers for qRT-PCR, NCBI Primers-BLAST online program was used. Information regarding the primers of these genes can be found in Supplementary Table 1. The 10 μL qRT-PCR reaction contained 1.0 μL of cDNA, 0.5 μL each primer at 10 μM, 5.0 μL of 2×SYBR^®^ Premix Ex Taq^™^ II (TaKaRa, Dalian, China), and 3 μL deionized water. The reaction was carried out on a LightCycler^®^ 96 Real-Time PCR system (Roche Applied Science, Indianapolis, USA). Internal control was the GAPDH gene while the 2^-ΔΔCT^ method was used to assess expression. The procedure for qRT-PCR amplification is shown: 95 °C for 3 min; 35 cycles at 95 °C for 30 s, 60 °C for 30 s, and 72 °C for 20 s; and a final extension for 10 min at 72 °C. The data were statistically analyzed using SPSS V 21.0 (SPSS Inc., Chicago, IL, USA). Measurement of one-way and repeated-analyses of variance followed by Dunnett’s test were carried out. The data are shown in the form of the means ± SE with significance set at p≤ 0.05.

### Protein extraction and iTRAQ reagent labeling

In the context of several experiments, an approach called isobaric tags for relative and absolute quantification (iTRAQ) has been successfully used. The samples were used for RNA-seq as well as iTRAQ analysis. For protein extraction, the lysis of 2 g each of the samples was done in lysis buffer 3 (TEAB with 1 mM PMSF, 8 M Urea, 10 mM DTT and 2 mM EDTA, pH 8.5) with 2 magnetic beads of 5 mm diameter. These were kept in a Tissue Lyser for 2 min at 50 Hz so that the proteins were released and subjected to centrifugation at 25,000 x g for 20 min at 4 °C, The transferring the supernatant into a fresh tube, 10 mM DTT (dithiothreitol) was added for reduction at 56 °C for 60 mins, followed by alkylation using 55 mM IAM (iodoacetamide) at room temperature, in dark for 45 min. This was then centrifuged at the same conditions described above.

The concentration and quality of proteins was assessed with a Bradford assay of the supernatant and confirmed with 12% SDS-PAGE.100 mM TEAB was used to dilute 100μg of protein solution with 8M urea that was subjected to digestion using Trypsin Gold (40:1: protein: enzyme, Promega, Madison, WI, USA) at 37°C overnight. Subsequently, an iTRAQ Reagent 8-plex Kit was used to label samples, followed by combining the differently labeled peptides, desalting using a Strata X C18 column (Phenomenex), finally vacuum drying in accordance to instructions of the manufacturer’s protocol. After this step, fractionation of these pooled mixtures was done on a Shimadzu LC-20AB high pressure liquid chromatography (HPLC) Pump system with a high pH RP column. This was followed by nanoelectrospray ionization and subsequent tandem mass spectrometry (MS/MS) on a Triple TOF 5600 system (SCIEX, Framingham, MA, USA). Triplicate experiments were done.

### Proteomics database processing

The ProteoWizard tool was utilized to convert raw MS/MS data into “.mgf” format followed by searching these exported files with Mascot (v 2.3.02) in this project against the database (*Gallus gallus 5.0, https://www.ncbi.nlm.nih.gov/assembly/GCF_000002315.4/*). Identification required the presence of minimum one unique peptide. Parameters set include: MS/MS ion search; Trypsin enzyme; 0.1 Da was the mass tolerance of fragment; Monoisotopic mass values; Variable modifications oxidation (M) and iTRAQ8plex (Y); 0.05 Da was tolerance for Peptide mass; Fixed modifications Carbamidomethyl (C), iTRAQ8plex (N-term), iTRAQ8plex (K); Database I-ZAwBa007 (50596 sequences); Database_info transcriptome. Identification was done using peptides that reached a confidence level of 95%.

The iTRAQ peptides were analyzed in a quantitative format with IQuant automated software [38]. This software incorporates a method based on machine learning called Mascot Percolator [39], that can rescore results from databases to yield a significant scale of standards. A 1% FDR was used to pre-filter PSMs that was used to measure the confidence level. A set of confident proteins was assembled using the parsimony principle also termed as the “simple principle”. Using the basis of picked protein FDR strategy [40], following protein inference with FDR at protein level lesser than or equal to 0.01, a FDR of 1% was measured in order to limit the number of false positives. The quantification of proteins involved identification of proteins, correction of impurities of tags, normalization of data, imputation of missing values, calculation of protein ratio, statistical approaches followed by presentation of results.

The proteins that had a p-value lower than 0.05 and a fold change of 1.2 were classified as DEPs. Analysis of metabolic pathways was in accordance to KEGG while analysis of other databases: COG and GO were in lieu of earlier research [41]. The hypergeometric test was applied to analyze DEP enrichment analysis in KEGG and GO. The equation was as follows:

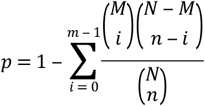

Where *N* represents the quantity of identified proteins that were linked to data from GO and KEGG analysis. *n* represents the measure of DEPs within *N. M* represents the quantity of proteins linked to a pathway or term of GO or KEGG, and *m* represents the quantity of DEPs linked to a pathway or term of GO or KEGG. An enrichment of differential proteins was considered when the p-value ≤ 0.05. The DEGs analysis was the same as the DEPs analysis.

### Transcriptomics and proteomics: Association analysis

Genes are regulated at multiple levels during the expression process. At present, most studies have reported that the expression consistency between mRNAs and their corresponding proteins is not very high, so a combined analysis of the proteome and transcriptome is helpful for discovering the regulation of gene expression [42]. Cluster 3.0 software was utilized to analyze analysis of clusters for DEP expression with that of transcripts in order to recognize transcripts and DEPs with similarity across various tissues with a graphical output on Java Treeview software. All expression data related to proteomics and transcriptomics were analyzed, and the Spearman correlation coefficient was calculated using R software [43]. Associations between the expression of mRNA and proteins from these respective “omics” were quantified followed by analysis for enrichment in GO as well as KEGG, in order to study the potential role of DEGs/DEPs in metabolism or signal transduction. The combined analysis parameters of transcriptomic and proteomic were given in Supplementary Table 2.

## Results

### Analysis of data from RNA sequencing

The Illumina HiSeq 4000 platform was employed to sequence 6 preparations of cDNA library. The data of RNA-seq shows high consistency among the libraries (Table 1). A total of 44.06 MB clean reads and 45.05 MB clean reads were obtained from the 6 cDNA libraries. Approximately 71% of the total reads could be mapped to the reference genome used for chicken while Q20/Q30 was more than 94%. The data is indicative of data of high quality sequencing that can ensure results of reliability for further investigation.

**Table 1.** Gene expression and clean reads analyses in the “Yufen I” H line feather follicle.

| Sample name | Raw reads/Mb | Clean reads/Mb | Clean Reads Q30/% | Mapping rate/% | Uniquely mapped rate/% | Total Gene Number | Total Transcript Number |
| --- | --- | --- | --- | --- | --- | --- | --- |
| A1 | 66.38 | 44.06 | 94.88 | 74.23 | 67.56 | 19054 | 36729 |
| A2 | 69.62 | 45.05 | 94.74 | 71.66 | 62.18 | 18640 | 33464 |
| A3 | 72.86 | 44.23 | 95.00 | 70.56 | 63.28 | 18669 | 33808 |
| B1 | 66.38 | 44.90 | 94.74 | 71.65 | 65.75 | 18567 | 33874 |
| B2 | 69.62 | 44.18 | 94.84 | 67.83 | 62.22 | 18565 | 34097 |
| B3 | 66.38 | 44.62 | 94.80 | 71.97 | 64.73 | 18793 | 33858 |

### Identification of DEGs

To identify genes involved in feather coloration, DEGs were detected in the “Yufen I” H line using PossionDis software [44]. Of total 21,306 identified genes, 597 novel genes and 12,687 novel isoforms were identified in chicken feather follicle libraries in this work. A total of 209 (83 upregulated and 126 downregulated) DEGs were detected at the two different locations of the feather follicle tissues using a less than 2 fold change and FDR ≤ 0.001. All of the DEGs are illustrated in Supplementary Table 3. Earlier research has shown several genes that involved in pigmentation (e.g., *ASIP, KITLG)* [45].

In order to corroborate whether the RNA-seq gene expression data were accurate and reproducible, 7 genes subjected to selection for qRT-PCR in the two groups. The expression of *KITLG, FZD10, WNT7B, WNT9A, WNT11, PVALB* and *MED23* was validated by qRT-PCR (Fig 2). As results from both these analyses were consistent with each other, the reliability of the sequencing is indicated.

**Fig 2.**
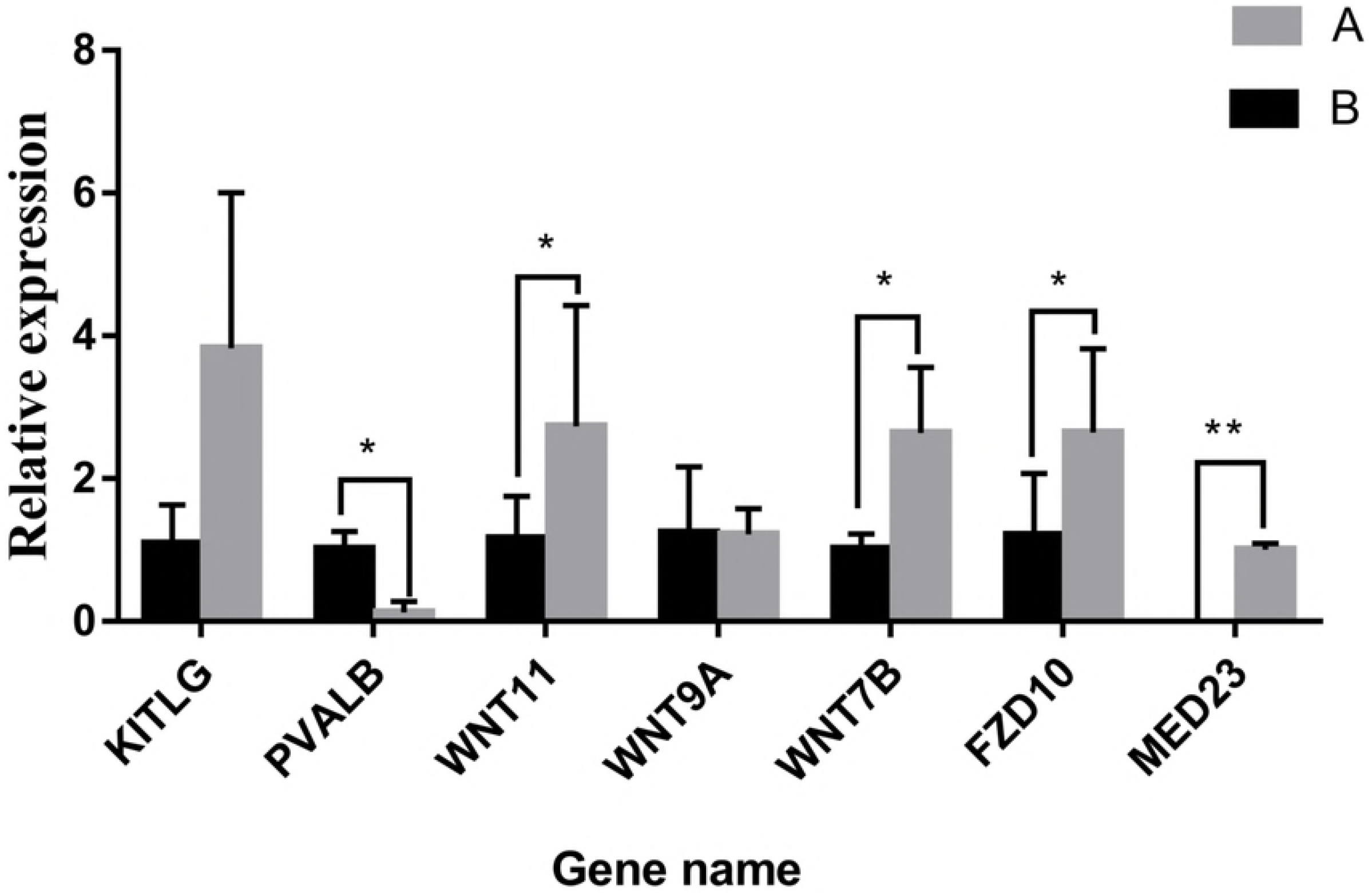
Verification of DEGs via qRT-PCR. The 2^-ΔΔCT^ method was used for data analysis and the housekeeping gene was GAPDH. Data shown on the vertical-axis represents the relative expression. The significant differences between two groups were determined by applying the unpaired Student’s t-test. * 0.01 < p < 0.05, ** p < 0.01. The presentation of all data is done as means ± standard error (SE).

The downy feathers of chickens are reportedly replaced by the second generation at the age of 6 weeks. At the age of 8 weeks, the second generation of feathers begins to molt and a large number of third-generation feathers begin to appear. At this time, the feather color tends to be stable. We collected feather follicles at 0, 2, 4, 6, 8, 10 and 12 weeks from the dorsal neck of the “Yufen I” H line chicken. Using qRT-PCR, we found that *MED23, FZD10, WNT7B* and *WNT11* gene expression peaked at approximately 8 weeks, which is consistent with the molting cycle (Fig 3). Furthermore, *FZD10, WNT7B and WNT11* are located in the melanogenesis pathway, and *MED23* specifically expressed in the dorsal feather follicles of the neck with black and white-striped feathers.

**Fig 3.**
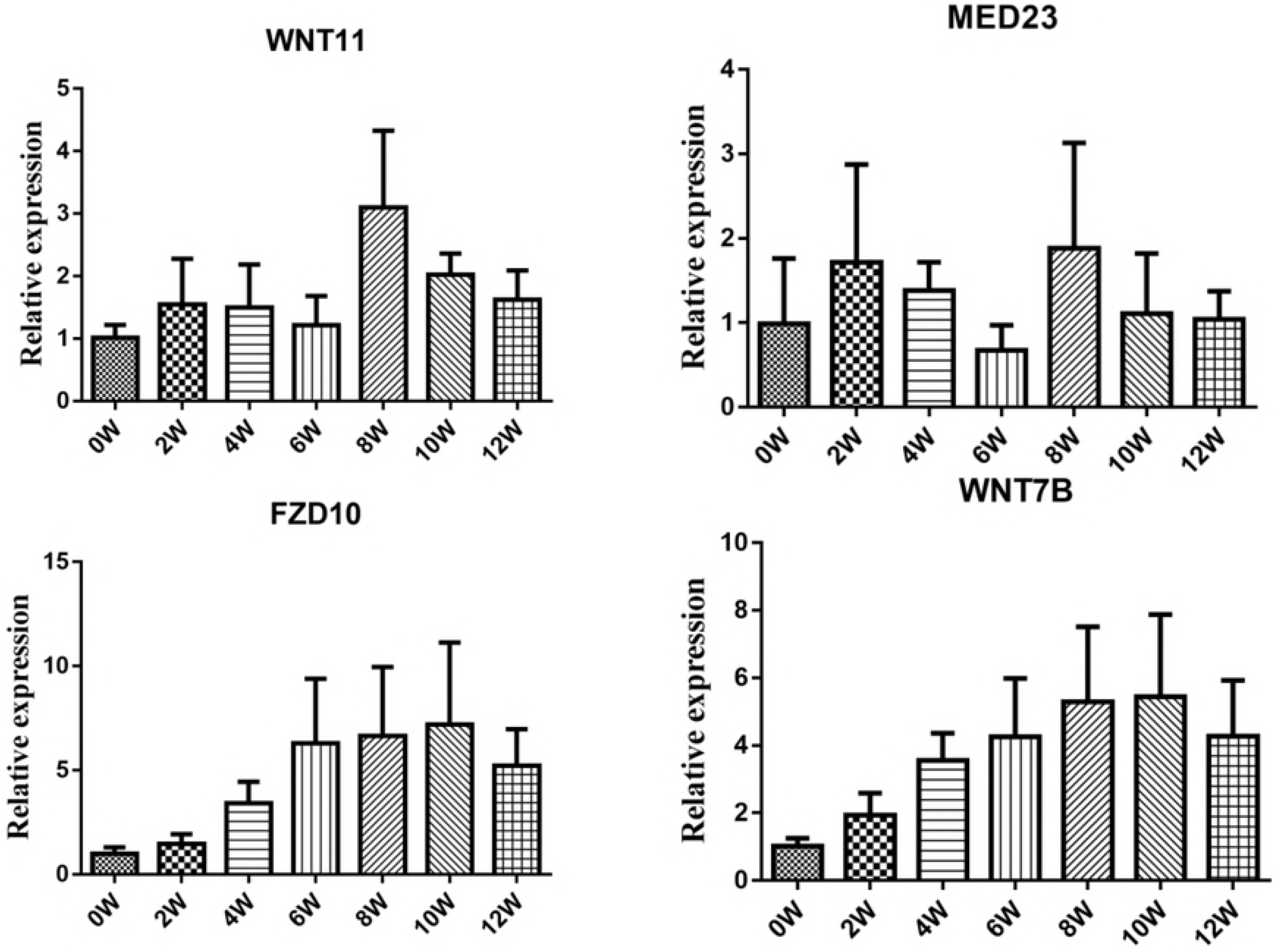
qRT-PCR validation of DEGs involved in feather cycle. Changes of the DEGs in feather follicles at 0, 2, 4, 6, 8, 10 and 12 weeks from the dorsal neck of the chicken.

### Metabolic pathways and GO analysis of DEGs

KEGG pathway and GO analyses were performed in order to interpret the exact roles played by the DEGs that regulated feather pigmentation. Using Blast2GO, we classified the DEGs with GO terms and KEGG pathways according to their functions. With the results of annotation from KEGG and GO, the official classification of DEGs was done followed by functional enrichment study in these databases using phyper, an R software function.

Classification of 209 DEGs with GO terms was done with the Blast2GO platform in terms of functional roles with an abundance of the following terms: “biological processes”, “molecular functions” and “cellular components” (FDR ≤ 0.01). Many genes were associated with cell part, cell, organelle, cellular process as well as binding. The categories with abundance were the processes associated with cell, single-organism, metabolic process and multicellular organisms, as well as regulation of biological process in the biological processes category. Cells, cell part and organelles showed maximum abundance in the category called cellular component. Majority of the unigenes could be classified into binding and catalytic activity functions in terms of function (Fig 4A).

**Fig 4.**
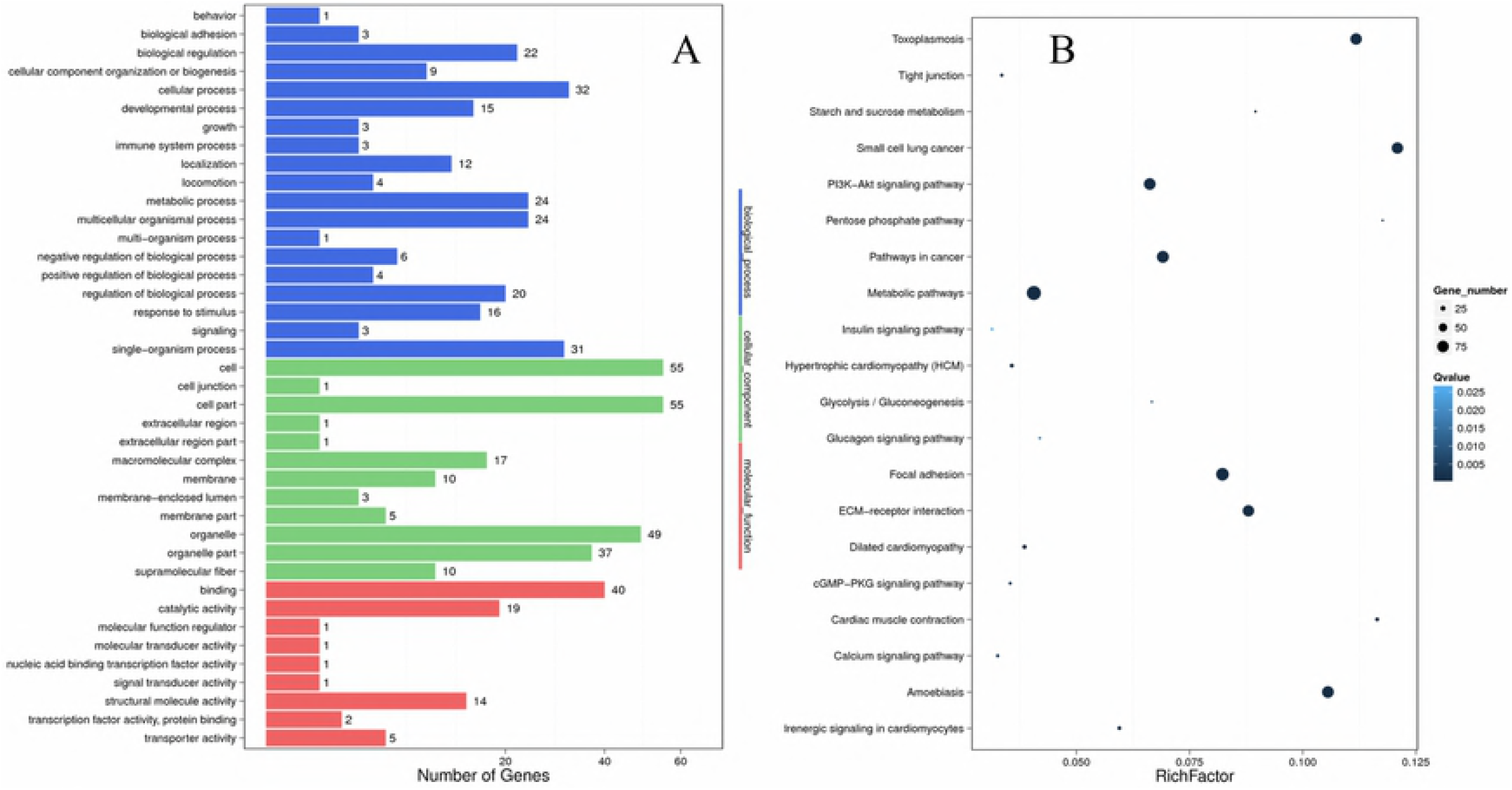
KEGG and GO analysis of DEGs. (A) GO functional annotation histogram of the DEGs. The vertical axis represents the three categories of GO, and the horizontal axis represents the gene number, and the number of genes considered as the differences in the proportion of the total. The GO annotations are classified in three different basic categories including cellular component, biological processes, and molecular function. (B) The degree of enrichment of the first 20 entries in the pathway. The pathway names are represented on the vertical axis, the horizontal axis represents the pathway corresponding rich factor. The ratio of the number of DEGs and all annotated genes in the pathway is defined as the rich factor.

As a result, mapping of 190 DEGs to 175 pathways in KEGG was done (FDR ≤ 0.01); therein, 188 DEGs were enriched in environmental information processing, of which 81 DEGs were enriched at the signaling molecules and interaction level and 107 DEGs showed enrichment at the level of signal transduction. The lowest Q value were found for the following 20 pathways: focal adhesion, metabolic pathways, signaling pathway of PI3K-Akt, cGMP-PKG, Calcium and that of adrenergic signaling in cardiomyocytes showed significant enrichment in two groups (Fig 4B).

### iTRAQ data analysis

iTRAQ analysis or proteomics was done to supplement data from transcriptomics for the same samples. The range of masses of identified protein was 10 to 100 kDa while the average coverage was challenged for groups more than 100 kDa. This data is indicative of reliable proteomic analyses [46]. This technique aided identification of 293,356 spectra in the samples that was subjected to data filtering to obtain 51,631 unique spectra that could be matched with 23,244 unique peptides, to totally identify 5,203 proteins (Table 2).

**Table 2.** Information of identified protein in the chicken feather follicle.

| Sample name | Total spectra | spectra | Unique Spectra | Peptide | Unique Peptide | Protein |
| --- | --- | --- | --- | --- | --- | --- |
| Yufen I | 293,356 | 60,886 | 51,631 | 25,545 | 23,244 | 5,203 |

### Functional classification and annotation of DEPs

The proteins expressed differentially were identified with the P-value < 0.05 and fold change ≥ 1.2 between the dorsal and ventral feather follicles of the neck. In brief, 382 DEPs (160 upregulated and 222 downregulated) were detected in the feather follicles. This was followed by classification into functions of COG categories. Clearly, changes were observed in the levels of proteins involved in functions such as general function prediction only; posttranslational modification, protein turnover; signal transduction mechanisms; translation, ribosomal structure and biogenesis; transcription and cytoskeleton (Fig 5A). DEPs were subjected to enrichment and clustering using functional analyses of GO and KEGG. In the GO function analysis, the cell part, intracellular part, and cytoplasm of cellular component; the binding, catalytic activity and ion binding of molecular function; and the biological regulation, response to stimulus and metabolic process of biological processes, were found to be different between the samples respectively (Fig 5B). Regarding KEGG pathway functions associated with the feather follicles, significant differences were found in the following signaling pathways: calcium, PI3K-Akt, mTOR, cAMP, MAPK, Wnt, cGMP-PKG, Jak-STAT and adrenergic signaling in cardiomyocytes, melanogenesis, tyrosine metabolism, and Melanoma. We displayed the top 20 enriched pathway in Fig 5C. At present, under feather regeneration treatment, in the dorsal follicles of the neck compared with the ventral follicles of the neck, segment polarity protein disheveled homolog DVL-3 isoform X6 protein (DVL3), 1-phosphatidylinositol 4,5-bisphosphate phosphodiesterase beta-1-like protein (LOC107052863), 1-phosphatidylinositol 4,5-bisphosphate phosphodiesterase beta-1 protein (PLCB1), GTPase KRas isoform X2 protein (KRAS), matrix-remodeling-associated protein 8 precursor protein (MXRA8), guanine nucleotide-binding protein G(q) subunit alpha protein (GNAQ), parvalbumin muscle isoform X1 protein (PVALB), calcium/calmodulin-dependent protein kinase type II subunit beta protein (CAMK2B) and calcium/calmodulin-dependent protein kinase type II subunit alpha isoform X3 protein (CAMK2A) were differentially expressed and mainly enriched for the melanogenesis pathway. The results indicated that upon defeathering stimulation, the DEPs related to feather pigmentation were mainly concentrated in the melanogenesis pathway.

**Fig 5.**
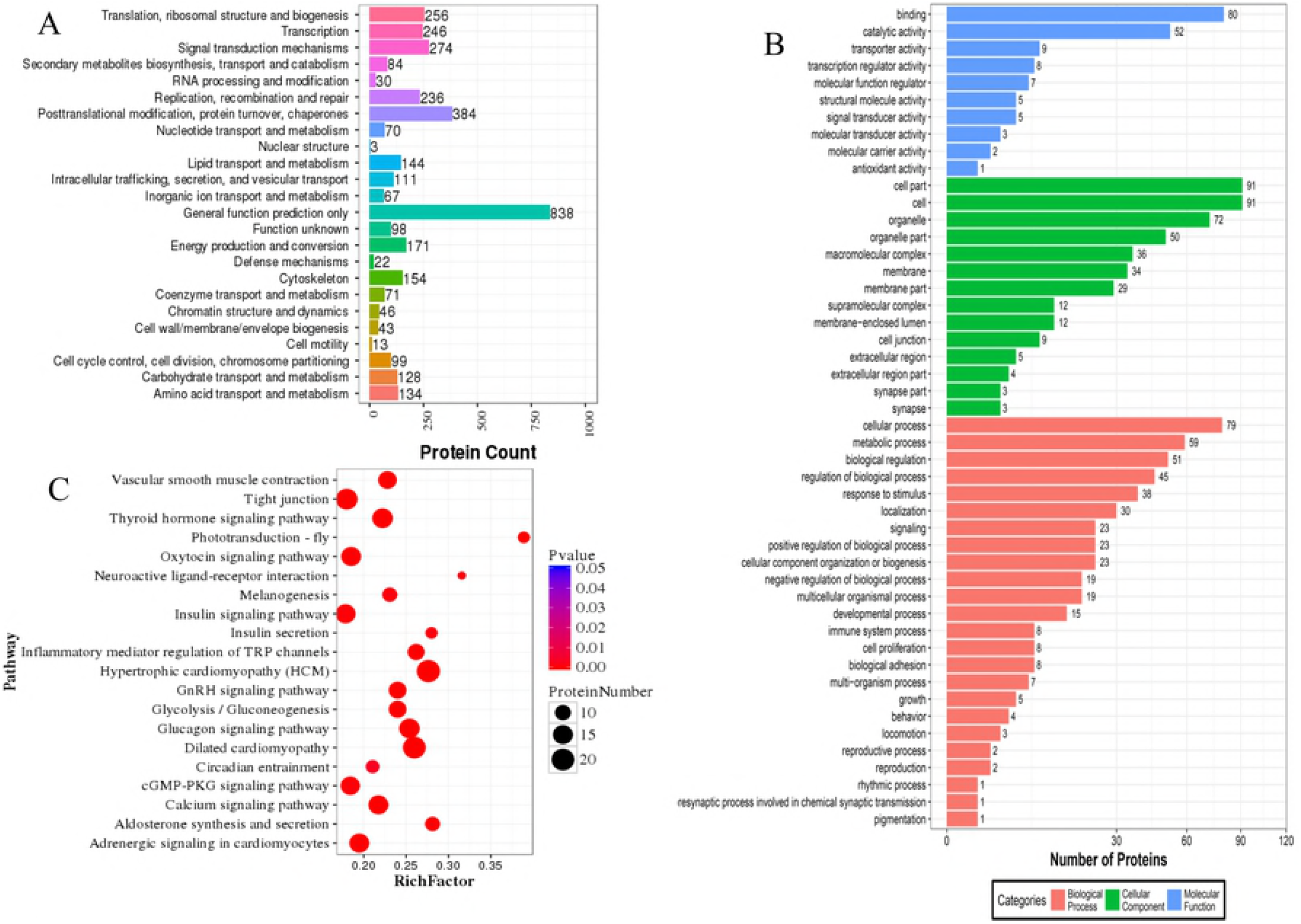
The DEPs analysis in terms of GOG, GO and KEGG. (A) The horizontal axis represents the COG term, the vertical axis represents the corresponding protein count illustrating the protein number of different function. (B) GO functional annotation histogram of the DEPs. The three categories of GO are presented on the vertical axis under the GO term, the horizontal axis represents the gene number, and the number of genes accounted for by differences in the proportion of the total. The GO annotations are classified in three different basic categories including cellular component, biological processes, and molecular function. (C) The name of the pathway is mentioned on the vertical axis, and the pathway matching the rich factor are mentioned on the horizontal axis. The rich factor refers to the ratio of the number of DEPs in the pathway and the number of all annotated proteins in the pathway. After testing multiple hypothesis, Q values were completed with corrected P value in the range of 0-0.05. The enrichment was considered significant if P value were closer to zero.

### Integrating transcriptomic and proteomic results

A majority of earlier reports are suggestive of a weak correlation between the expression of mRNA with protein attributed to posttranscriptional regulation or posttranslational modification or errors in experiments [47–49]. Nevertheless, the flow of information from RNA to proteins is the crux of the central dogma [50,51]. To allow for genes that are expressed differentially in terms of the transcriptome and proteome as well as singling out vital genes, we performed an integration of DEGs and DEPs. For multi-omics data, GO terms and KEGG pathways were enriched the levels of transcriptome and proteome, respectively. Then, the data from the two groups were integrated and analyzed, which was conducive to the study of gene expression regulation at the level of coexpression of gene sets [52–55].

To compare the proteomic with the transcriptomic, we compared the 382 DEPs with the 209 DEGs. According to the analysis results, only 49 genes meeting the criteria overlapped (Supplementary Table 5). The changes at the level of transcripts and proteins showed a weak correlation for the proteins that were quantified. Biological pathways were elucidated with a change in the reports from statistics for protein levels when the changes in mRNA were zero. To investigate the overall correlation between these transcripts and differentially expressed proteins, matching of all identified mRNAs with DEPs was done followed by transformation of DEP and transcript volume ratios into log2 forms. An investigation of changes at both transcript and protein levels revealed a weak correlation (Spearman correlation coefficient, R= 0.0840) for all the genes and proteins assessed. We then calculated the correlation between the 382 DEPs and the 209 DEGs, and a positive correlation of R = 0.3006 was obtained when all significantly changed proteins with a cognate mRNA were considered (Fig 6). This result showed a modest correlation between the levels of mRNA and protein (between the proteome and transcriptome), this was consistent with previously reported results [56], and accounted for the complexity of the gene expression regulation mechanism [42,57–59].

**Fig 6.**
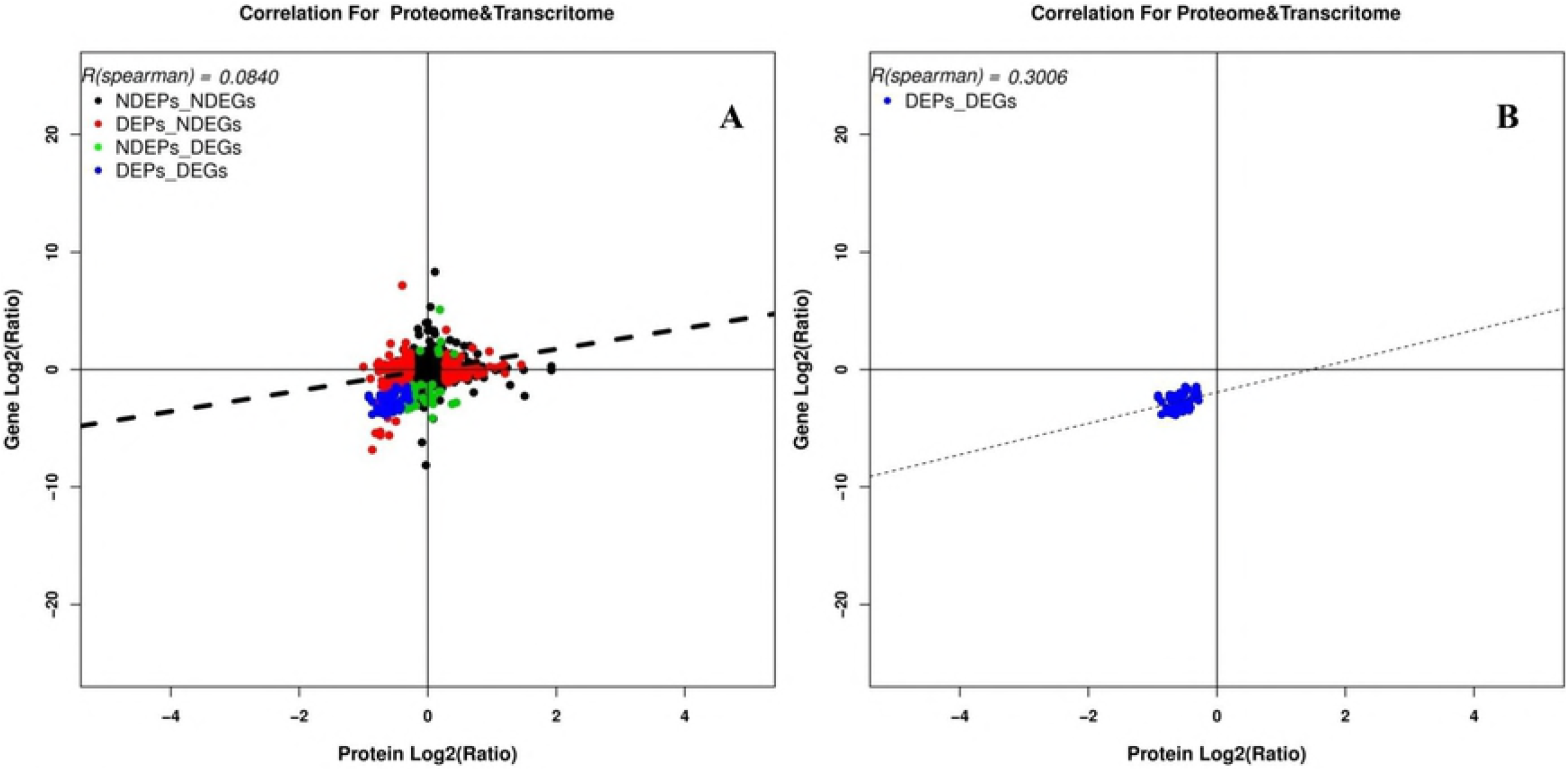
Correlations between the expression of proteins and genes. The vertical-axis represents the protein expression level, the horizontal-axis represents the genes expression level. (A) Scatter plots of the correlation between data sets of genes evaluated in both proteomic and gene transcripts analyses. (B) Scatter plots and coefficients of DEPs and DEGs correlation.

49 DEGs/DEPs were assigned GO terms to assess their functions that encompassed a vast range of cellular components, molecular functions and biological processes (Supplement Fig 1). The GO analysis indicated that the DEGs/DEPs relevant to biological processes termed cellular, metabolic and biological regulation are possibly related to our study and showed that the DEGs/DEPs were relevant to molecular functions including catalysis or binding or structure.

Assignment to COG functional categories indicated that excluding carbohydrate transport and metabolism and the cytoskeleton, the DEGs/DEPs were classified into the categories of inorganic ion transport and metabolism and general function prediction.

Additionally, among the components of a single pathway the extent of comity between proteins and mRNA was studied. In total, 382 DEPs and 209 DEGs were mapped to 166 biological pathways, of which 12 pathways showed significant enrichment in both groups. This study involved the determination of vital DEPs and DEGs, involved in melanogenesis and the following signaling pathway: calcium, cGMP-PKG and adrenergic signaling in cardiomyocytes using pathway enrichment analysis (Fig 7).

**Fig 7.**
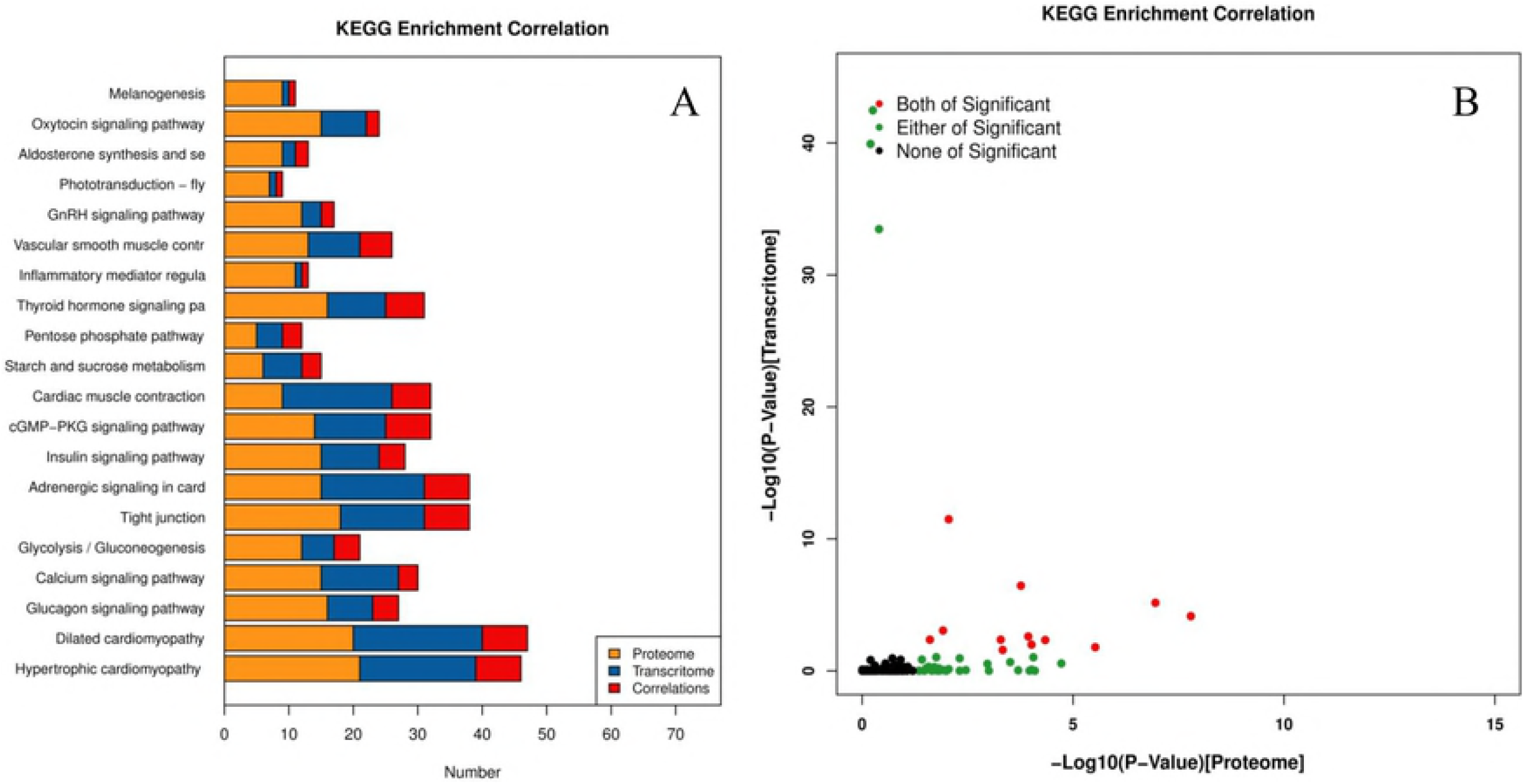
Correlation of KEGG enrichment between transcriptome and proteome. (A) Number of KEGG enrichment correlation transcriptome and proteome. (B) The overview scatter diagram of KEGG enrichment correlation between the transcript level and protein level of genes.

## Discussion

Feathers have evolved to have diverse coloration that exhibit a wide spectrum of colors arranged in noticeable patterns. Either physical or chemical or a combination can form these colors. Melanin is the major factor responsible for the different types of colors described in the existing literature.

### Feather regeneration and pigmentation after feather removal

Similarly to hairs, the remarkable property of molting and regeneration cycles is seen in feathers [60]. During the regenerative process for feathers and feather follicles, melanocytes and melanoblasts were activated by reactive stimuli; furthermore, melanocyte stem cells are potential sources of new melanocytes. The location of these stem cells include the bases of the follicle and epidermal collar as well as the dermal papilla apex [61,62]. While melanin formation in the skin is an ongoing process, this pathway in the case of hair is functional throughout the phase of growth. The process is not functional in the catagen stage (a phase of transition where renewal of a follicle occurs at the termination of anagen) as well as telogen stage [63]. In the event of a loss of a feather (intentional or accidental), an old follicle generates a new one in a span of 2 weeks [64]. In our study, when we pulled out the feather and stimulated the feather follicle to develop again, new feathers were formed. Moreover, black and white-striped feathers and white feathers regrew in the dorsal and ventral neck areas, respectively.

### DEGs and DEPs related to melanin synthesis

Pigmentation based on melanin is linked to a sequence of genes with initiation and regulation of production of the pigment involving many signaling pathways as well as transcription factors such as the tyrosine kinase receptor *KIT* with *SCF* ligand, and *MITF* [65]. Many genes in melanogenesis pathway are related to *MITF,* this is a sole transcription factor of the microphthalmia family that is involved in regulation of melanocytes. *MITF* target genes regulate melanocyte pigmentation [66]. Previous studies revealed that *MITF* was involved in the development of melanocytes, along with reports of color of plumages of Japanese quail and chicken and mutation in this gene [67].

In this study, a total of 8 DEGs (*ASIP, KITLG, FZD10, WNT7B, WNT9A, WNT9B, WNT11, PVALB*) and 9 DEPs (DVL3, KRAS, MXRA8, GNAQ, PVALB, CAMK2B, CAMK2A, PLCB1, LOC107052863) were found to be involved in the melanogenesis pathway (Fig 8). *MED23* was specially expressed and significantly upregulated in dorsal follicles of the neck. A new study revealed that zebrafish pigmentation is regulated by *MED23* via a modulation of the function of *MITF* as an enhancer [68]. GNAQ was significantly upregulated in the dorsal follicles of the neck in this pathway. Researches showed that the *GNAQ* activates *MITF* via MAPK pathway, and then affected melanin synthesis [69]. As a consequence, melanin synthesis can be regulated via influencing the expression of the *MITF* gene and then regulates the development of feather coloration. In addition, we found that *MED23, FZD10, WNT7B* and *WNT11* genes expression peaked at approximately 8 weeks in the “Yufen I” H line, which is consistent with the molting cycle. The results are indicative of an extensive and vital role of these genes in melanin synthesis, which is consistent with previous research results. Therefore, we speculated that the above genes were related to the formation of Columbian plumage in “Yufen I” H line.

**Fig 8.**
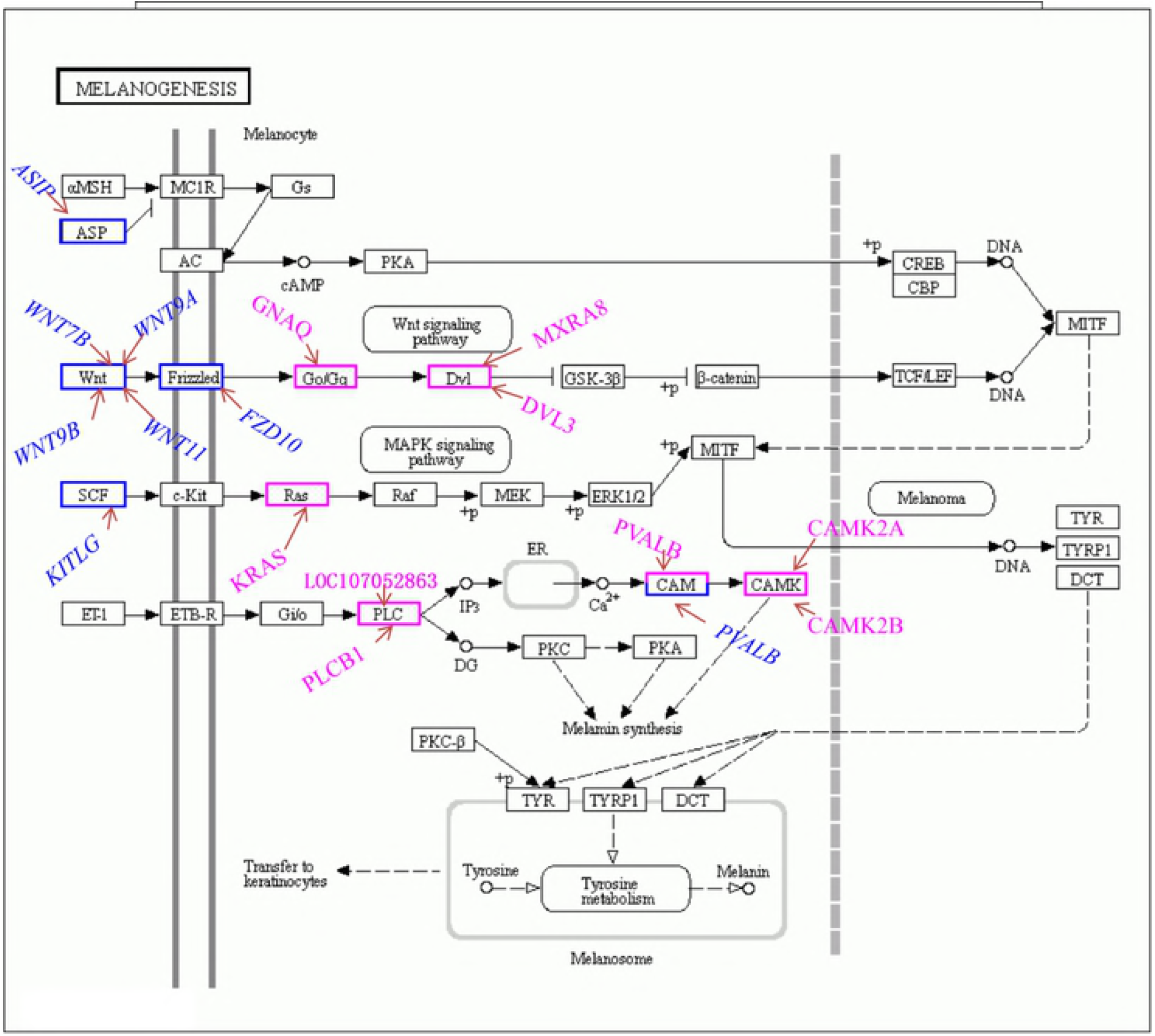
Differentially expressed feather color genes and proteins identified in the analyzed chicken feather follicles and their involvement in the melanogenesis pathway. The DEGs with a blue frame, the DEPs with a pink frame.

### Pathways related to melanin synthesis

To further understand the DEGs and DEPs in the pathways that regulate chicken feather color development, we performed KEGG pathway analysis of DEGs and DEPs (p < 0.05). Correlation analysis and integration were performed on DEPs and DEGs annotated in the same pathway. Combining these results with those of the transcriptome and proteome analyses showed that, melanogenesis, adrenergic signaling in cardiomyocytes, calcium and cGMP-PKG signaling pathway were highly prevalent in comparisons of the different feather color occurrence locations of “Yufen I” H line feather follicles. Therefore, we speculate that these four pathways may be related to the Columbian plumage of “Yufen I” H line, although we primarily focused on the melanogenesis pathway in this study. We found that the DEGs were mainly concentrated in the upstream of the pathway, while the DEPs were concentrated in the downstream of the pathway (Fig 8).

Interestingly, 5 proteins (PLCB1, LOC107052863, PVALB, CAMK2A, CAMK2B) and the *PAVLB* gene regulated the melamin synthesis in melanogenesis pathway. Few studies have previously reported on the correlation between melamine and pigmentation. The effect of melamine on the skin color of darkbarbel catfish showed that the factors necessary for melanin formation may be suppressed by intake of melamine, resulting in a significant reduction in melanin in the skin of the dorsal surface [70]. Therefore, we speculate the 5 genes that regulated melamine synthesis may also be related to feather color deposition by affecting melanin synthesis.

## Conclusion

In summary, an incorporated and strong approach of analyzing the transcriptome and proteome was utilized to study the working of melanin synthesis in feathers. Strikingly, this original report of a transcriptomics and proteomics analysis of the feather color in chicken follicles describes and reveals a set of differentially expressed genes which are putatively involved in color of feathers along with other physiological functions. The expression of *MED23, FZD10, WNT7B, WNT11* and GNAQ were significantly different in terms of feather follicles with black and white stripes versus white feather color, and elucidating the functional roles of these genes in the regulation of feather color will be of particular interest in future studies. These results provide a potential understanding of the working of Columbian plumage in the “Yufen I” H line and solid genetic resources that allow for selection of birds uniform for plumage to allow for breeding.

## Acknowledgments

This study was carried out with the aid provided by the Agriculture Research System of China (CARS-40-K04), Program for Innovation Research Team of Ministry of Education (IRT16R23), Key Science and Technology Research Project of Henan Province (151100110800).

## Data availability statement

All raw data have been deposited under Accession number: SRR7973871 to NCBI Sequence Read Archive.

---

**Supplement Fig 1. GO enrichment correlation between transcriptome and proteome.** (A) The number of GO enrichment correlation between transcriptome and proteome. (B) The scatter diagram overview of GO enrichment correlation between the level of transcript and protein of genes.

**Supplementary Table 1. Information regarding the specific primers used for the qRT-PCR.**

**Supplementary Table 2. The combined analysis parameters of transcriptomic and proteomic.**

**Supplementary Table 3. Differentially expressed genes between the dorsal feather follicles of neck and the ventral feather follicles of neck.**

**Supplementary Table 4. Differentially expressed proteins between the dorsal feather follicles of neck and the ventral feather follicles of neck.**

**Supplementary Table 5. Correlation DEPS and DEGs.**

